# Bile is a route of excretion for homogentisic acid in alkaptonuria

**DOI:** 10.1101/2023.12.18.572165

**Authors:** BP Norman, H Sutherland, PJM Wilson, DA Rutland, AM Milan, AT Hughes, AS Davison, M Khedr, JC Jarvis, JA Gallagher, G Bou-Gharios, LR Ranganath

**Author notes:** **Corresponding author:** Brendan P Norman, Address: Institute of Life Course & Medical Sciences, University of Liverpool, William Henry Duncan Building, 6 West Derby Street, Liverpool, UK. L7 8TX.

## Abstract

Altered activity of specific enzymes in phenylalanine-tyrosine (phe-tyr) metabolism results in incomplete breakdown of various metabolite substrates in this pathway. Increased biofluid concentration and tissue accumulation of the phe-tyr pathway metabolite homogentisic acid (HGA) is central to pathophysiology in the inherited disorder alkaptonuria (AKU). Accumulation of the metabolites upstream of HGA, including tyrosine, occurs in patients on nitisinone, a licenced drug for AKU and hereditary tyrosinaemia type-1, which inhibits the enzyme responsible for HGA production. The aim of this study was to investigate the phe-tyr metabolite content of key biofluids and tissues in AKU mice on and off nitisinone to gain new insights into the biodistribution of metabolites in these altered metabolic states. The data show for the first time that HGA is present in the bile in AKU (mean [±SD] = 1003[±410] μmol/L; nitisinone-treated AKU mean [±SD] = 45[±23] μmol/L). Biliary tyrosine, HPPA and HPLA are also increased on nitisinone. Urine was confirmed as the dominant elimination route of HGA in untreated AKU, but with indication of biliary excretion and possible metabolism of HGA by the gut microbiome. These data provide new insights into the pathways of phe-tyr metabolite biodistribution and metabolism, showing for the first time that hepatobiliary excretion contributes to the total pool of metabolites in this pathway. Our data suggest that biliary elimination of organic acids and other metabolites may play an underappreciated role in disorders of metabolism.

**Take-home message:** This paper presents the first observation of elevated hepatobiliary circulation of metabolites associated with disease in alkaptonuria, including homogentisic acid in addition to tyrosine and the tyrosine metabolites 4-hydroxyphenylpyruvic acid and 4-hydroxyphenyllactic acid on nitisinone treatment.

## Introduction

The phenylalanine-tyrosine (phe-tyr) pathway is the major metabolic route through which excess phenylalanine and tyrosine is catabolised. Phe-tyr metabolism predominantly occurs in the liver and involves the conversion of these amino acids through several intermediate steps, resulting in the metabolite products fumarate and acetoacetate. Mutations in genes encoding the various enzymes of the phe-tyr pathway have deleterious consequences, resulting in the accumulation of metabolites that directly cause a range of specific and potentially fatal diseases (Figure S1).^1^

Alkaptonuria (AKU; OMIM #203500) is an inherited disorder of the phe-tyr pathway, in which mutations in the gene encoding the homogentisate 1,2-dioxygenase enzyme (HGD; E.C.1.12.11.5) result in its inability to metabolise the intermediate homogentisic acid (HGA).^2^ Despite maximal urinary excretion of HGA in untreated AKU, HGA concentration remains elevated in plasma and accumulates in tissues throughout the body.^3^ HGA accumulation in tissues gives rise to a pigment formed from oxidation of HGA, a slowly progressing process called ochronosis. Ochronosis is central to the chronic multisystemic features of AKU, which include cardiac valve disease, premature arthritis, bone fractures and rupture of ligament, tendon and muscle.^4–6^

In recent years, nitisinone (2-(2-nitro-4-(trifluoromethyl)benzoyl)cyclohexane-1,3-dione) has been established as an effective agent for treatment of AKU. Nitisinone reduces plasma and urinary HGA, slows progression of disease in patients^7,8^ and completely arrests ochronosis in AKU mice.^9,10^ Nitisinone is the only known disease modifying agent for AKU and hereditary tyrosinaemia type 1 (HT-1) but its inhibition of HGA production causes the accumulation of metabolites upstream of HGA in phe-tyr metabolism, including tyrosine, 4-hydroxyphenylpyruvic acid (HPPA) and 4-hydroxyphenyllactic acid (HPLA). Accumulation of tyrosine is a major health concern in nitisinone-treated AKU and HT-1.^11–16^

Ochronosis is well established as the central cause of disease pathogenesis in AKU, but there remains much to learn about this metabolic pathway. It is clear that ochronotic pigment is an oxidation product of HGA, but the exact chemical structure of the pigment remains unknown.^17,18^ The molecular mechanism through which HGA-derived pigment becomes bound to cartilaginous tissue, the preferential site of pigment accumulation, is also not fully understood.^19–21^ Metabolite data from the ‘Suitability of Nitisinone in AKU’ (SONIA 1^22^ and SONIA 2^7^) clinical trials have provided insights into the metabolism of HGA, including identification of the major HGA biotransformation pathways,^23^ and have enabled inferences on the daily biodistribution rates of HGA in plasma and total body water compared with urinary excretion.^24,25^ It is not currently possible to directly measure all the HGA-derived products in the ochronotic pathway, but the blockade of HGA production by nitisinone in these studies has been studied as a way to unmask the amount of flux that would otherwise be directed down this pathway in untreated AKU. SONIA 1 and SONIA 2 both showed a marked increase in the summed concentrations of phe-tyr pathway metabolites in total body water, but not in urine, in patients on nitisinone treatment. The interpretation of this finding was that approximately half of the total daily HGA produced is excreted in the urine. It is possible that the remaining half of the daily HGA accumulates in tissues as HGA and/or pigment, but estimates of lifetime accumulation based on this rate of deposition seem implausible. An alternative, but as yet unconfirmed, explanation for the total phe-tyr flux unaccounted for in untreated AKU is that biliary secretion is a significant elimination route of HGA from the liver.^24^

The enterohepatic circulation is the circulation of bile and its constituents between the liver and gastrointestinal tract. It is well known that endogenous substrates and xenobiotics, including drugs and metabolites are excreted into bile from hepatocytes.^26^ The liver is the major site of phe-tyr metabolism^27^ and the transporter machinery for small molecule anions such as HGA exists at the canalicular surface of hepatocytes for secretion into the bile, namely transmembrane transporter multidrug resistance protein 2 (MRP2).^28,29^ Once secreted into bile, it is proposed that HGA is secreted into the small intestine, from which both secondary absorption and faecal excretion are possible. We tested the hypothesis that bile HGA is a contributor to the total phe-tyr metabolite pool in AKU. Phe-tyr metabolites have not been directly measured in bile in patients with AKU due to the invasiveness of accessing this biofluid. The Hgd^-/-^ mouse accurately recapitulates the AKU metabolic state and the early stages of disease pathophysiology including tissue ochronosis.^27^ This mouse model of AKU provides an opportunity to systematically investigate bile metabolite composition both in untreated AKU and in response to nitisinone treatment under controlled conditions.

## Methods

### Chemicals and reagents

Deionised water was purified in-house by DIRECT-Q 3UV water purification system (Millipore, Watford, UK). Methanol, isopropanol (Sigma–Aldrich, Poole, UK), formic acid (Biosolve, Valkenswaard, Netherlands) and ammonium formate (Fisher Scientific, Schwerte, Germany) were LC/MS grade. Nitisinone, perchloric acid and H_2_SO_4_ were from Sigma-Aldrich.

### Animal details

All mice studied were on a C57BL/6 genetic background. AKU mice were generated from targeted knockout of Hgd (Hgd^-/-^).^27^ For AKU mice, the nitisinone-treated and no treatment groups each comprised four mice (2 male, 2 female). Mean age (±SD) was 13.5(±0.9) weeks in the nitisinone group and 12.5(±1.6) weeks in the no treatment group. Wild type mice were three females all aged 29.1 weeks. For bile measurements, samples were taken from an additional two AKU untreated mice; 1 male and 1 female, both aged 9 weeks.

All mice were housed, maintained and experimental procedures carried out within the University of Liverpool Biological Services Unit in specific pathogen-free conditions, under project licence PP8132802, in accordance with UK Home Office guidelines, under the Animals (Scientific Procedures) Act 1986. Mice were culled by Schedule 1 approved methods and tissues harvested without delay. All mice were housed under pathogen-free conditions, in cages of up to five mice, with 12-hr light/dark cycle, and food and water available *ad libitum*. Mice were drug/test-naïve at baseline. In the nitisinone-treated group, nitisinone was added to all drinking water to a final concentration of 4 mg/L.

### Sample collection and analysis

Blood samples were collected from venous tail bleeds into lithium heparin microvettes (CB 300 LH, Sarstedt, UK) and kept on ice until processed. Whole blood was centrifuged at 1500x *g* for 10 min at 4 °C and plasma supernatant acidified with 10% v/v 5.8 M perchloric acid then vortexed. Acidified plasma samples were centrifuged at 1500x *g* for 10 min at 4 °C and the supernatant removed.

Urine was sampled on a single-collection basis into 0.5 mL microcentrifuge tubes. Urine samples were acidified by addition of 5% v/v 1 M H_2_SO_4_.

Post schedule 1 cull, the thyroid and gall bladder were excised. Bile was removed from the gall bladder. Faecal pellets were freshly collected. All samples were snap frozen in liquid nitrogen. All tissues and biofluid samples were stored at -80 °C prior to analysis.

Whole thyroid tissues, which were pooled among groups to provide sufficient material, and individual faeces samples were thawed on ice in 1.5 mL microcentrifuge tubes prior to metabolite extraction. Extraction was performed by addition of 50% of the final volume of 0.58 M perchloric acid to samples and homogenised in tubes by manual handheld homogeniser. The homogeniser tip was then washed in the second 50% volume of perchloric acid, and the wash transferred to the tissue homogenate. Tissue extract concentrations (tissue weight/extraction buffer volume) were 250 mg/mL for pooled thyroid and 125 mg/mL for faeces. All tissue homogenates, urine and bile samples were centrifuged at 1500x *g* for 10 min at 4 °C, supernatant was removed and diluted 1:1000 sample:internal standard-containing diluent as with plasma and urine samples. Plasma and urine samples were prepared and quantified for phe-tyr pathway metabolites by liquid chromatography triple-quadrupole tandem mass spectrometry (LC-QQQ-MS) following published plasma (serum) and urine protocols.^30–32^ Bile, faeces and thyroid samples were analysed using the aforementioned serum phe-tyr protocol.

### Data processing and statistical analysis

Raw LC/MS data files were processed in MassHunter QQQ Quantitative Analysis software (v. B.09.00; Agilent) following published protocols.^30–32^ Statistical analyses were performed in Prism (v. 6.01; Graphpad). Statistical significance tests employed were paired t-tests for paired data (plasma, urine and faeces) and Kruskal-Wallis tests for unpaired data (bile). P values <0.05 were considered significant.

Summed values for the total phe-tyr metabolite pool measured here were calculated for each plasma, urine and bile sample. The combined raw μmol/L concentrations of all phe-tyr metabolites were calculated and summed values derived for each biofluid or tissue sample as follows. Summed plasma μmol/L concentrations were multiplied by 60% of body weight (g)^33^ to reflect total body water concentrations. Total 24-hr urine metabolite output was estimated by multiplying the summed μmol/L urine concentrations by published 24-hr urine output volume (in L/100 g body weight) in C57BL/6 mice, which equates to 7.1 mL/100 g body weight in males and 9.9 mL/100 g body weight in females.^34^ The gall bladder bile pool was also estimated using published data on total gall bladder bile volume in C57BL/6 mice; 4 μL bile per g liver weight.^35^ Liver weight estimates were calculated as 4.3% of total body weight in males and 4.11% of total body weight in females (C57BL/6 mean organ weight data available from the Jackson Laboratory: https://www.jax.org/jax-mice-and-services). Summed bile μmol/L concentrations were then multiplied by the estimated total gall bladder bile volume (L) to derive total gall bladder metabolites.

## Results

### Individual phe-tyr pathway metabolites

For plasma, urine and faeces, metabolite concentrations were compared between paired samples taken from the same AKU mice at baseline then at one week on nitisinone or no treatment. Terminal bile metabolites were compared between three groups of mice: untreated AKU mice, AKU mice at one week on nitisinone and untreated wild type mice. Faeces concentrations are expressed as μmol/mg tissue. A summary of measured metabolite concentrations is shown in Table S1.

Terminal bile HGA concentration showed a marked increase in untreated AKU compared with both nitisinone-treated AKU and wild type (Figure 1). There was a difference >20-fold between untreated AKU (mean [±SD] = 1003[±410] μmol/L) and nitisinone-treated AKU (mean = 45[±23] μmol/L). All bile HGA concentrations in wild type mice fell below the lower limit of quantification (LLOQ; 3.1 μmol/L). Further analysis within the untreated AKU group revealed a clear sex difference in bile HGA, with a statistically significant increase in male mice (P = 0.018). Analysis of the other phe-tyr pathway bile metabolites showed clear increases in tyrosine, HPPA and HPLA in nitisinone-treated AKU mice (Figure 2), and nitisinone-treated males also had increased concentrations of these metabolites (Figure S7). Bile phenylalanine was also increased in untreated AKU; statistically significant difference compared with treated AKU.

**Figure 1.**
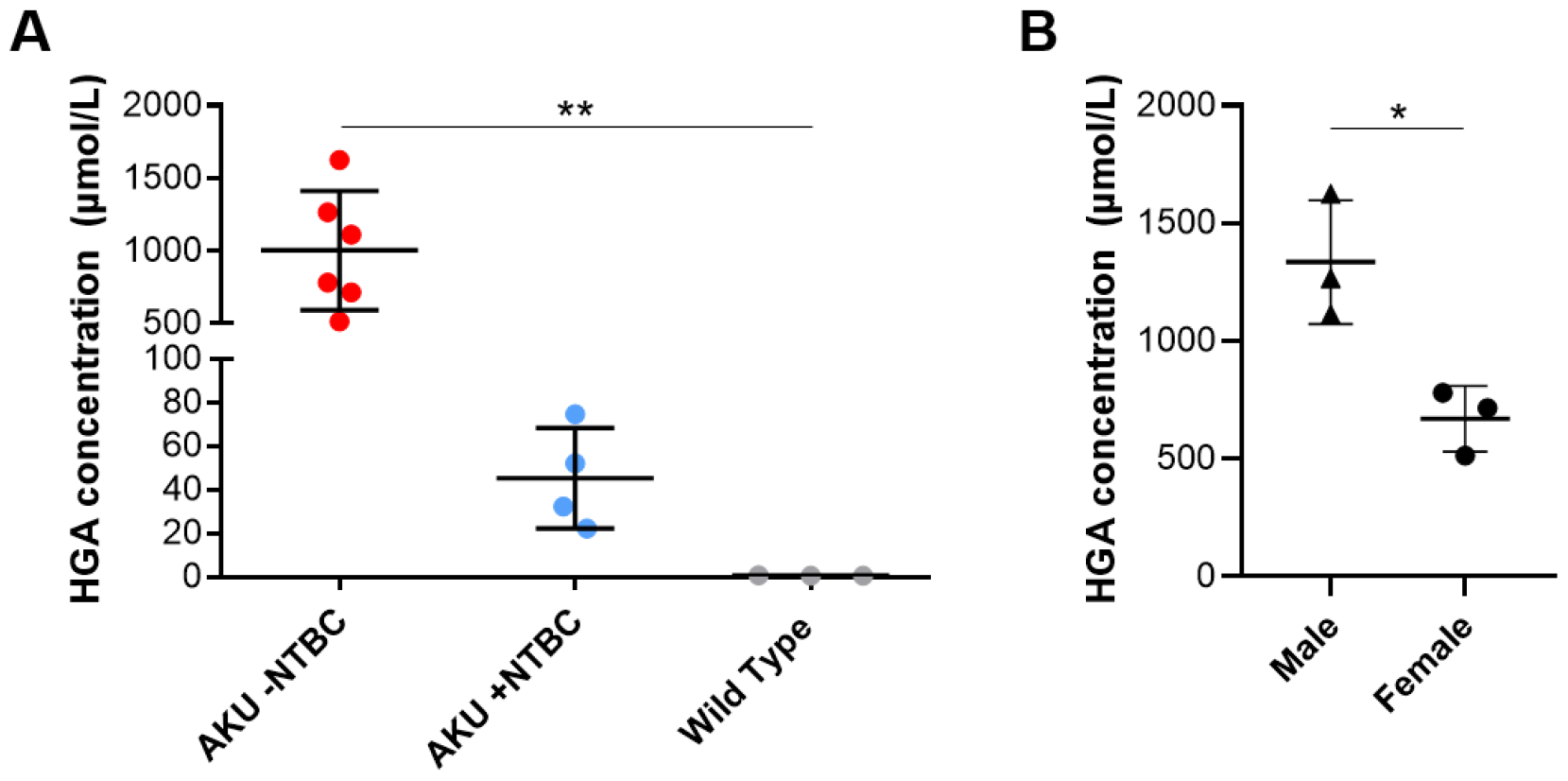
Bile HGA is increased in AKU. Bile HGA showed a marked increase in Hgd^-/-^ untreated AKU mice compared with AKU mice at 7 days on nitisinone (NTBC) and wild type mice (A). Bile HGA is increased in male versus female untreated Hgd^-/-^ mice (B). *P <0.05, **P <0.01 (B).

**Figure 2.**
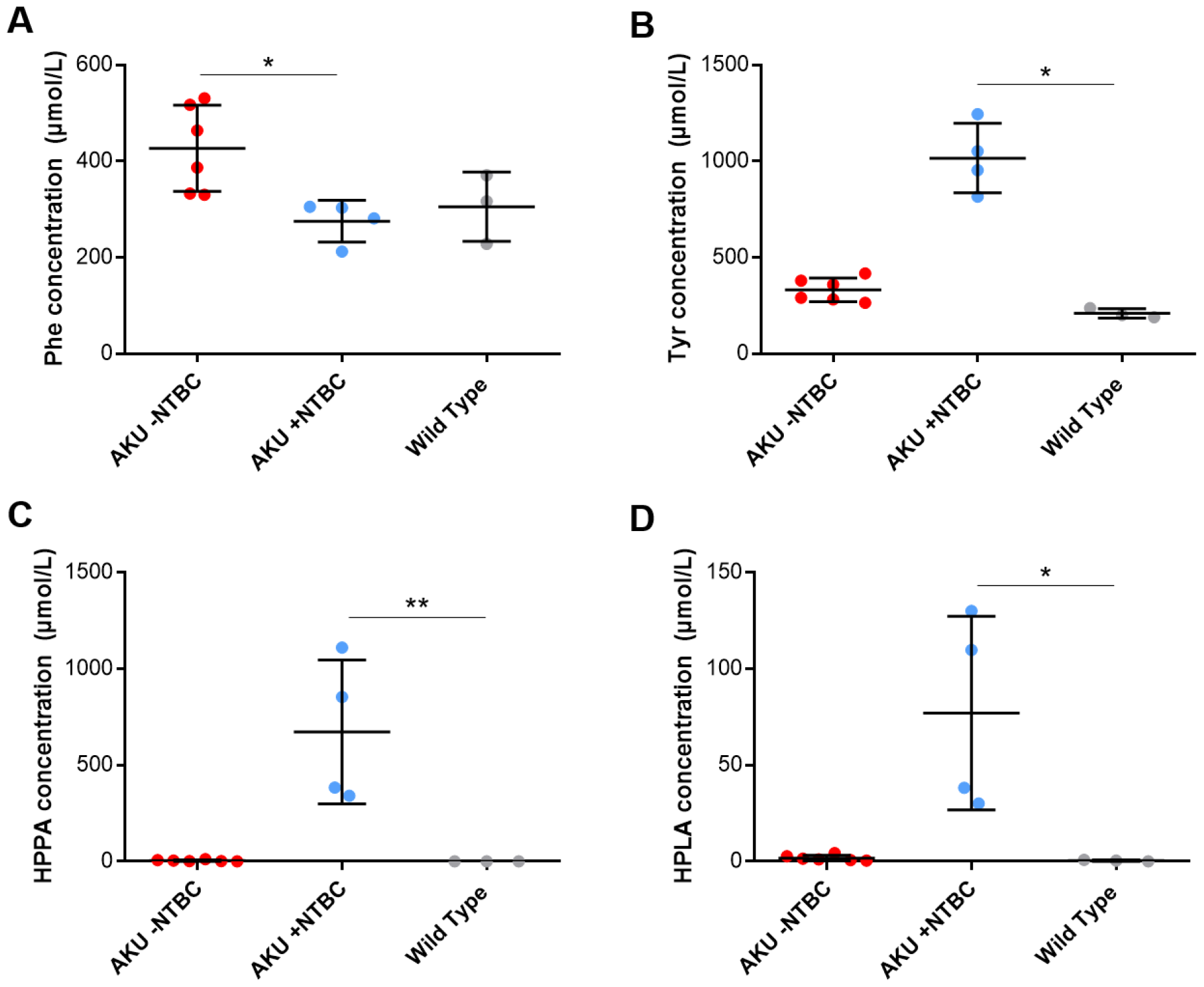
Bile phe-tyr pathway metabolites in AKU and their response to nitisinone. Statistically significant differences were observed for each of the phe-tyr pathway metabolites analysed beside HGA. Phenylalanine was increased in untreated Hgd^-/-^ versus Hgd^-/-^ at seven days on nitisinone (NTBC) (A). Tyrosine, HPPA and HPLA were markedly increased in Hgd^-/-^ on nitisinone (B-D). *P <0.05, **P <0.01.

Plasma and urine showed marked decreases in HGA, as expected, at one week on nitisinone in AKU mice (Figure S2). Mean plasma HGA at baseline was 211(±58) μmol/L then 8(±3) μmol/L on nitisinone; P <0.01. Mean urine HGA was 136.7(±38.2) mmol/L at baseline then 9.4(±2.3) mmol/L on nitisinone; P <0.01. Plasma HGA decreased at one week in the untreated control group but this difference was not statistically significant; P = 0.076. Urine HGA showed no clear change at one week in untreated controls; P = 0.66. Faeces HGA data were more variable with no clear trend across serial samples in treated and untreated groups. There were two outlier faeces samples with HGA at 1971 μmol (at baseline) and 268 μmol (at one week on nitisinone) per g tissue, but the other samples remained close to the LLOQ for HGA.

Plasma and urine showed the expected clear increases in tyrosine, HPPA and HPLA on nitisinone treatment. Signals for these metabolites, in addition to phenylalanine, in faecal extracts were more variable. Faecal phenylalanine and tyrosine were above the LLOQ but no statistically significant differences were observed from baseline in treated or untreated groups, despite a decrease in these metabolites in 3/4 mice at one week on nitisinone. HPPA and HPLA were not consistently detected across samples, except for one mouse on nitisinone (Figures S3-S6).

Data from thyroid homogenates pooled within the same three groups of mice showed clear group differences generally consistent with those observed for bile (Table S1). Thyroid HGA was detected only in the untreated AKU group, and tyrosine and HPLA were greater in nitisinone-treated AKU compared with untreated AKU and wild type.

### Summed phe-tyr pathway metabolites

Summed data were analysed for the paired samples taken at baseline then at one week on nitisinone/no treatment based on the approach previously described in patients with AKU^24,25^ (Figure 3). In line with previous reports in patients, a clear increase in summed plasma metabolites from baseline was observed at one week on nitisinone (P <0.01) but not at one week of no treatment (P = 0.14). For the one-week samples, summed plasma + bile was also increased on nitisinone treatment compared with no treatment (P <0.001). This increase in summed metabolites on nitisinone was no longer observed when urine data were considered; no difference was observed from baseline for summed urine metabolites, summed urine + plasma, or summed urine + plasma + bile metabolites.

**Figure 3.**
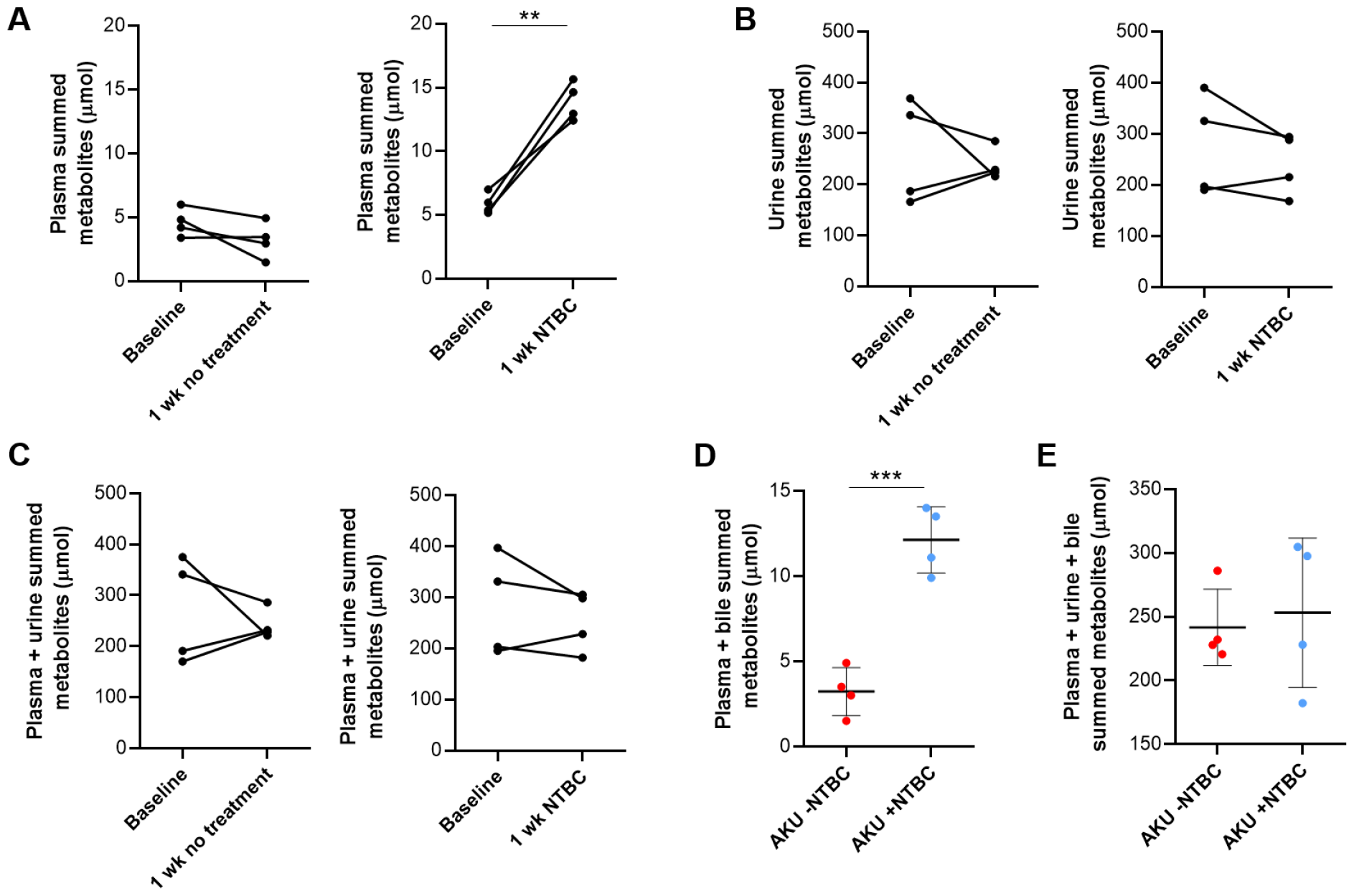
Summed phe-tyr metabolite data in treated versus untreated AKU. Summed phe-tyr metabolite data (phe, tyr, HPPA, HPLA and HGA) in AKU Hgd^-/-^ mice. A-C shows paired data; the same mice at baseline then at seven days on nitisinone (NTBC) or no treatment. D-E shows unpaired data incorporating bile measurements. A: plasma summed, B: urine summed, C: plasma summed + urine summed, D: plasma summed + bile summed, E: plasma summed + urine summed + bile summed. **P <0.01, ***P <0.001.

## Discussion

Untreated AKU and nitisinone-treated AKU result in two contrasting altered metabolic states characterised by differential accumulation of phe-tyr pathway metabolites. In untreated AKU, HGA accumulation and subsequent ochronotic pathway activity occurs due to oxidation of HGA, the metabolite that directly causes multisystem disease in this condition. Nitsinone inhibits the 4-hydroxyphenylpyruvate dioxygenase enzyme, which blocks the formation of HGA and ochronosis, but with invariable accumulation of metabolites upstream of HGA; phenylalanine, tyrosine, HPPA and HPLA. To our knowledge, this study is the first systematic demonstration that bile is a contributor to total pathway flux in both untreated and treated metabolic states. Our findings of biliary concentrations of approximately 1 mmol/L for HGA in untreated AKU and tyrosine in treated AKU indicate that bile is an important excretory route for the accumulating phe-tyr pathway metabolites in these conditions. To our knowledge, this is the first direct observation of biliary excretion of HPPA and HPLA, the metabolites of tyrosine, which also accumulate in patients on nitisinone. Our observations in mice provide new fundamental insights into the tissue and biofluid distribution of HGA. We confirm that urine is the major excretion route of HGA but show for the first time that HGA is also excreted via the biliary system and alimentary tract. Hepatobiliary excretion is referred to as ‘phase III’ detoxification of molecules,^36^ which adds to our previous observations that HGA undergoes detoxification by phase I and II metabolism in AKU.^23,37^ It is worth noting that our data may underestimate HGA biliary excretion, as previously reported HGA biotransformation products may also be excreted with HGA from the liver and therefore contribute to the total HGA pool.

Bile is a complex aqueous secretion uniquely formed in the liver, which is the major site of phe-tyr metabolism. Bile is approximately 95% water containing many constituents including bile salts, bilirubin, phospholipids, cholesterol, amino acids, steroids and hormones in addition to endogenous drugs and xenobiotics. Hepatobiliary secretion serves as an excretory route for a diverse range of substrates including xenobiotics and endogenous small molecules.^26^ The literature on biliary excretion of phe-tyr metabolites in the disorders affecting this pathway is scarce. A case report of a Danish patient with alkaptonuria suggests potential secretion of HGA in bile. This patient had recurrent gallstones over a 30-year period, and upon starting a low phenylalanine and tyrosine diet, bile duct stone formation immediately subsided, as did associated abdominal pain. The remission on the low phenylalanine and tyrosine diet strongly supports a role for HGA in the formation of bile stones.^38^ Compared with HGA, the circulation between hepatobiliary secretions and the intestine for amino acids is better understood.^39^ Amino acids including phenylalanine and tyrosine from biliary secretions undergo subsequent reabsorption throughout the intestine via the solute carrier transporter SLC6A19.^40^ Our finding of generally low or undetectable HGA in faeces indicates that HGA secreted in bile is reabsorbed in the intestine and/or metabolised by the gut microbiome. There is evidence that HGA is absorbed from the intestine when given orally to humans.^41^ Phenylalanine and tyrosine catabolism also occurs widely in microorganisms and various bacterial strains are known to express HGD.^42^ HGA is a central intermediate in bacteria, in which it has received considerable interest as a source of production for the pigment pyomelanin in multiple species.^17,43^

From the HGA concentrations established here, it is possible to estimate rates of daily HGA output via bile, faeces and urine in untreated mice using the published total 24-hour outputs of these biofluids/tissues in mice (Table 1). These calculations show the dominant urinary route of HGA output, resulting in 13.4 - 115.7 mg HGA/24-hours. Bile flow rate is reportedly greater in female mice,^44^ and bile HGA concentration was increased in males in this study, but the range of estimated 24-hour HGA output remains an order of magnitude lower than for urinary HGA output. Estimated bile output was more consistent across untreated AKU mice compared with faecal output due to the variability in faecal HGA concentrations. Two outlier faecal samples had increased HGA at 1971 μmol/g tissue and 268 μmol/g tissue, giving 24-hour faecal HGA output estimates of 0.1 - 0.01 mg from these samples. The remainder of faecal samples produced minimal 24-hour faecal HGA output estimates due to measured HGA concentrations below or close to the assay LLOQ. The reason for variability in faecal HGA is not clear, but potential factors include individual differences in a) metabolism of HGA by the gut microbiome and b) reabsorption efficiency of the HGA secreted into the intestine from bile. The source of the phe-tyr metabolites observed in faeces is also unclear, as they could be derived from hepatobiliary secretion, diet or microbial metabolism.

**Table 1.**
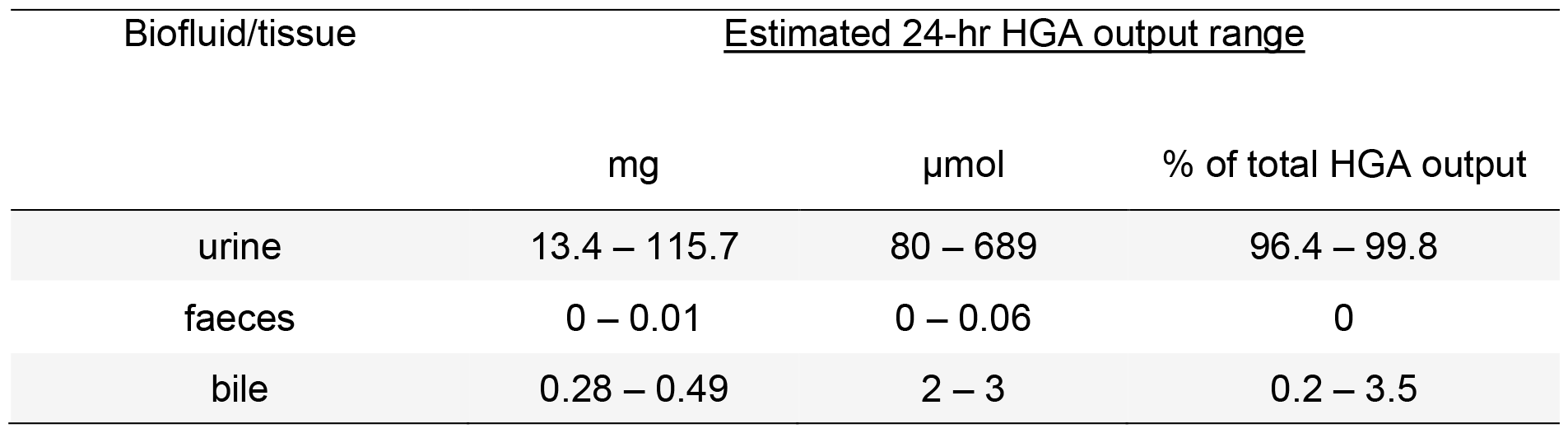
Estimated range of 24-hr HGA output in urine, faeces and bile in untreated AKU Hgd^-/-^ mice. Values represent 24-hr output of HGA calculated from measured tissue/biofluid concentrations and literature outputs of bile,^44^ faeces and urine.^45^ For urine, the range estimations are based on the lowest measured HGA concentration in untreated Hgd^-/-^ mice combined with the lower limit 24-hr urine output (0.85 mL); upper range calculated from the greatest measured urine HGA concentration combined with the upper limit 24-hr urine output (3.375 mL). 24-hr faecal output is based on estimated 14.5 mg total output of faeces per g of body weight.^45^ 24-hr bile output is based on the mean bile flow rate reported in mice, taking into account established sex differences in this measure: 2.5 mL/24-hr for females and 1.8 mL/24-hr for males.^44^

| Biofluid/tissue | <u>Estimated 24-hr HGA output range</u> |  |  |
| --- | --- | --- | --- |
|  | mg | μmol | % of total HGA output |
| urine | 13.4 – 115.7 | 80 – 689 | 96.4 – 99.8 |
| faeces | 0 – 0.01 | 0 – 0.06 | 0 |
| bile | 0.28 – 0.49 | 2 – 3 | 0.2 – 3.5 |

The data from summed phe-tyr metabolites for each biofluid suggest important differences in phe-tyr metabolism between mice and humans (Figure 3). While summed plasma metabolites showed the clear increase in nitisinone-treated subjects that has been characterised in multiple AKU patient cohorts,^24,25^ summed plasma and urine metabolites combined showed no clear difference in mice on *versus* off nitisinone. Increases on nitisinone were only observed for plasma summed metabolites and plasma + bile summed metabolites, suggesting that in AKU mice, consideration of urinary metabolite clearance negates the increase in summed plasma phe-tyr metabolites on nitisinone. This finding is in contrast to reports in humans that summed concentrations of the same metabolites in total body water combined with urine were approximately double in nitisinone-treated subjects; this effect was sustained over the 48-month duration of the SONIA 2 clinical trial.^7^ We propose that the reason for this difference between species is decreased activity in mice of the ochronotic pigment pathway derived from HGA, meaning that there is not the same degree of ‘unmasking’ of this pathway when on nitisinone compared with humans. In patients, the major contributor to increased summed metabolites is the elevated tyrosine, with minor contributions from the elevated HPPA and HPLA. Mice have nitisinone-induced tyrosinaemia to a similar degree as humans, and our data suggest that the major metabolic difference between the two species in this context is greater urinary excretion of HGA in AKU mice. Mean urinary HGA concentration (random urine) was 133.7 mmol/L in mice in this study, which is approximately 3.8 times greater than the concentrations of 35 mmol/L (24-hr collection urine) reported in SONIA 2. Such increased urinary HGA excretion is a possible reason for the observation that mice do not develop such extensive macroscopic tissue ochronosis and joint destruction as in human AKU, although other factors have been proposed including the shorter lifespan of mice and endogenous vitamin C production as an additional defence against HGA autoxidation.^9,46^ Renal elimination of HGA has not yet been studied systematically in mice but the importance of the kidney to HGA detoxification in patients is well known; the renal HGA elimination rate exceeds the theoretical renal plasma flow. Organic anion transporters appear crucial to this process, as the contribution of net tubular secretion of HGA is greater than glomerular filtration.^3^

We report the first direct observation of HGA accumulation in the thyroid in AKU, with concomitant thyroid tyrosine accumulation on nitisinone. The concentrations of HGA and tyrosine in thyroid are several orders of magnitude greater than their respective plasma concentrations when comparing the equivalent concentrations per weight of tissue/biofluid; tissue concentrations are in μmol per g of tissue, biofluid concentrations expressed in μmol per L of fluid (Table S1). Increased thyroid HGA is consistent with previous reports of ochronotic thyrocytes in AKU, indicated by Schmorl’s staining of patient thyroid tissue.^47^ Primary hypothyroidism has increased prevalence in AKU, in which thyroid gland dysfunction appears to be mainly attributable to exposure to HGA and/or HGA-derived pigment, over tyrosine. In a recent analysis, it was observed that for all cases of hypothyroidism reported in nitisinone-treated AKU patients, the hypothyroidism preceded the administration of nitisinone.^47^ Future investigations are required to establish the reason for thyroid as an apparent preferential site of tissue HGA accumulation.

Clear sex differences were observed in bile metabolite concentrations. HGA was higher in untreated male AKU mice, and tyrosine and the organic acids HPLA and HPPA were higher in males at one week on nitisinone (Figure S7). This finding may reflect greater phe-tyr pathway flux in males due to higher dietary intake and is consistent with greater AKU disease severity in male patients.^5^ The reason for higher bile phenylalanine in untreated AKU mice is not clear and requires further investigation; this finding is inconsistent with previous reports of increased plasma phenylalanine on nitisinone (vs untreated baseline) in AKU mice.^23^

Limitations of this study include those relating to one-off sample collections. The ‘snapshot’ concentrations obtained from biofluid/tissue samples enabled valuable estimations of total metabolite flux and output over 24-hr, but we acknowledge that studying the mice in metabolic cages would have enabled a more detailed readout of pathway flux and rates of specific metabolite elimination. Metabolic cages were avoided for several reasons, including the difficulty in acidifying urine for HGA stabilisation in this approach. Metabolic cages are also known to induce stress in rodents, which can affect the evaluation of important factors in phe-tyr metabolism including feeding habits and renal function.^45^ Constituents of bile, such as bile acids, show diurnal variation in mammals. In mice, this pattern is in phase with the fasting/feeding cycle, which results in release of gall bladder bile to the intestine and reabsorption to the liver during night-time feeding hours. Gall bladder bile acid concentration peaks during daytime hours in mice,^48^ therefore the daytime gall bladder bile metabolite concentrations reported here may represent values towards the maximal diurnal range. It was not possible to measure such diurnal bile metabolite fluctuations without much more invasive experimental procedures.

In conclusion, our observations provide new insights into the biodistribution of phe-tyr pathway metabolites in the altered metabolic states of alkaptonuria and tyrosinaemia induced by nitisinone. For the first time, we have shown that there is biliary secretion of each of the metabolites that accumulate in these conditions, including amino acids and organic acids. Urinary excretion is confirmed as the dominant route of HGA elimination in untreated AKU, but the findings highlight that bile and tissue metabolite content should be considered in attempts to model or quantify total pathway flux in AKU and also in disorders of metabolism more broadly.

## Author contributions

Conceptualisation, BPN, JAG, GB, LRR; Data curation, BPN; Formal analysis, BPN, AMM, ATH, ASD; Funding acquisition, BPN; Investigation, BPN, HS, PJMW, DAR; Resources, JCJ, GB, LRR; Writing – original draft, BPN; Writing – review and editing, all authors. All authors have read and agreed to the published version of the manuscript.

## Ethics approval

Research on laboratory animals reported in the manuscript was approved under project licence PP8132802 and performed in accordance with UK Home Office guidelines, under the Animals (Scientific Procedures) Act 1986.

## Patient consent statement

N/A

## Data sharing statement

Raw data reported in the manuscript will be shared by the corresponding author upon reasonable request.

