## Supplemental for "Bile is a route of excretion for homogentisic acid in alkaptonuria"

|  | Phe | Tyr | HPPA | HPLA | HGA |
| --- | --- | --- | --- | --- | --- |
| Plasma |  |  |  |  |  |
| AKU -NTBC | 74 (13) | 76 (27) | ND | ND | 83 (49) |
| AKU +NTBC | 75 (10) | 686 (53) | 64 (14) | 34 (10) | 8 (3) |
| Urine |  |  |  |  |  |
| AKU -NTBC | 10 (1.6) | 36 (8) | 1307 (556) | 250 (215) | 125,781 (9440) |
| AKU +NTBC | 21 (16) | 489 (356) | 102,426 (30,236) | 10,814 (7922) | 9350 (2346) |
| Wild type | 23 (11) | 91 (47) | 1264 (924) | 47 (17) | 6 (4) |
| Gall bladder bile |  |  |  |  |  |
| AKU -NTBC | 427 (90) | 332 (62) | 4 (4) | ND | 1003 (410) |
| AKU +NTBC | 275 (44) | 1016 (181) | 671 (373) | 77 (50) | 45 (23) |
| Wild type | 305 (72) | 209 (25) | ND | ND | ND |
| Faeces |  |  |  |  |  |
| AKU -NTBC | 44 (22) | 137 (58) | ND | ND | ND |
| AKU +NTBC | 40 (22) | 102 (53) | ND* | ND* | ND* |
| Wild type | 15 (4) | 47 (6) | ND | ND | ND |
| Thyroid |  |  |  |  |  |
| AKU -NTBC | 115 | 135 | ND | ND | 28 |
| AKU +NTBC | 104 | 556 | ND | 34 | ND |
| Wild type | 85 | 105 | ND | ND | ND |

**Table S1.** Summary of measured metabolite concentrations (mean [±SD]). Data from AKU mice are at one week of nitisinone treatment (AKU +NTBC) or no treatment (AKU -NTBC). WT mice were untreated. Biofluid concentrations are in µmol/L. Thyroid and faeces concentrations are µmol per g of tissue. Thyroid metabolite data were single measurements from pooled whole thyroid homogenates (n = 3 per pool).

* Mean values excluding one outlier sample in AKU +NTBC group with increased faecal metabolites (faeces concentrations in µmol per g of tissue: HPPA = 1194; HPLA = 558; HGA = 268).

ND: not detected. Measurement below assay lower limit of detection.

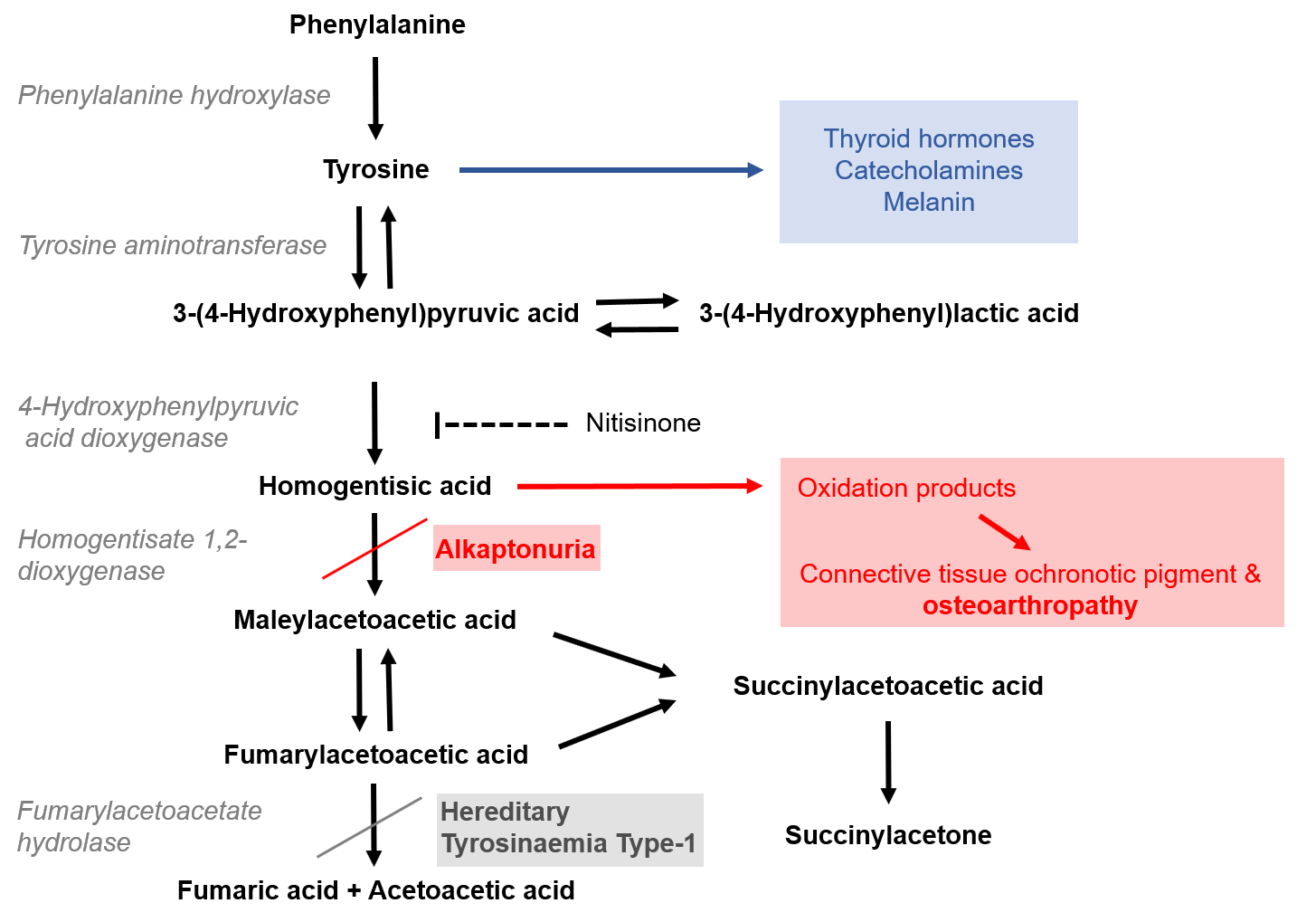

**Figure S1**. The phenylalanine-tyrosine metabolic pathway, with the enzyme defects in alkaptonuria (AKU) and hereditary tyrosinaemia type-1 (HT-1) highlighted, in addition to the inhibition of 4-hydroxyphenylpyruvic acid dioxygenase by nitisinone.

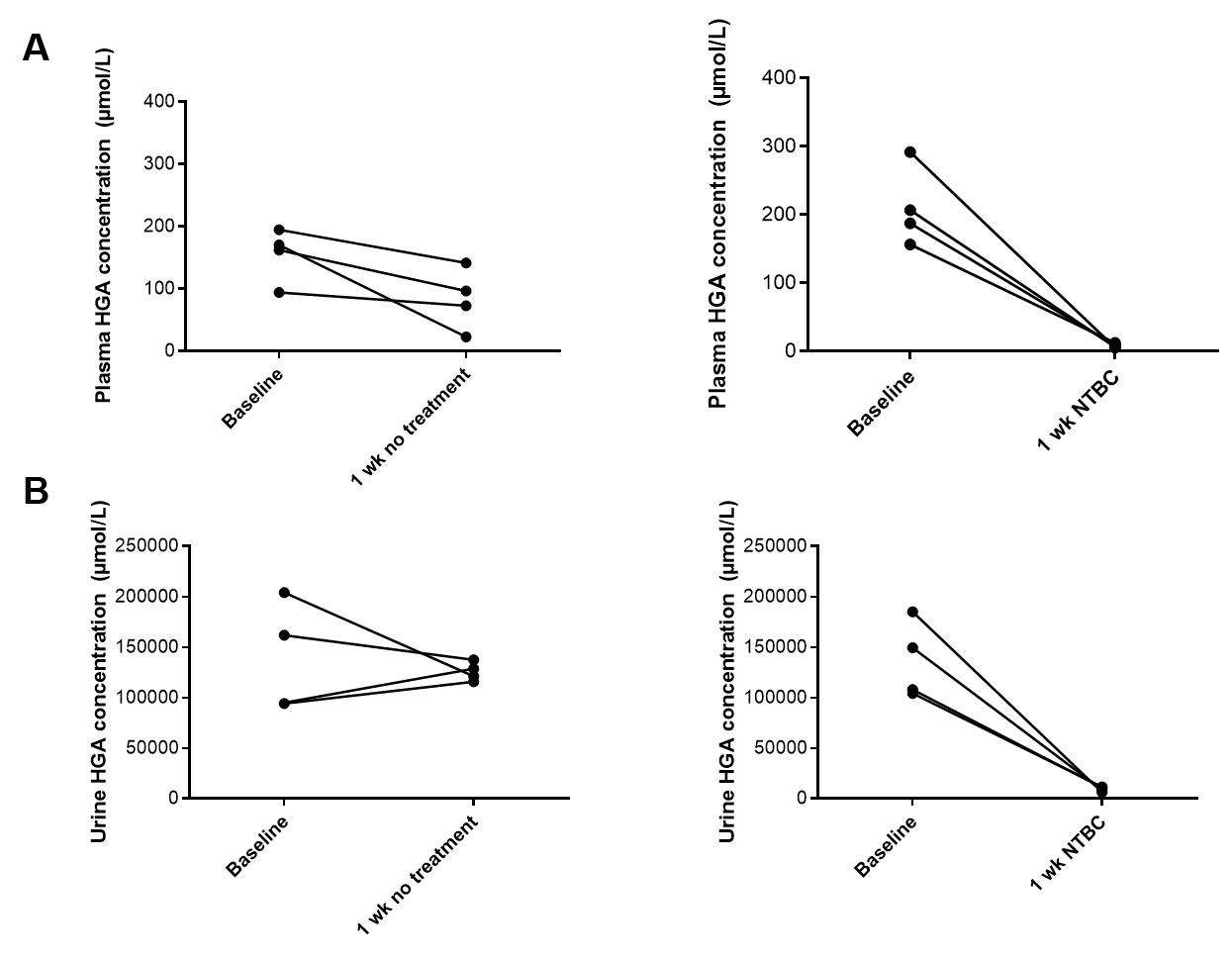

**Figure S2**. HGA concentrations at baseline then at one week on nitisinone (NTBC) or no treatment in plasma (A) and urine (B).

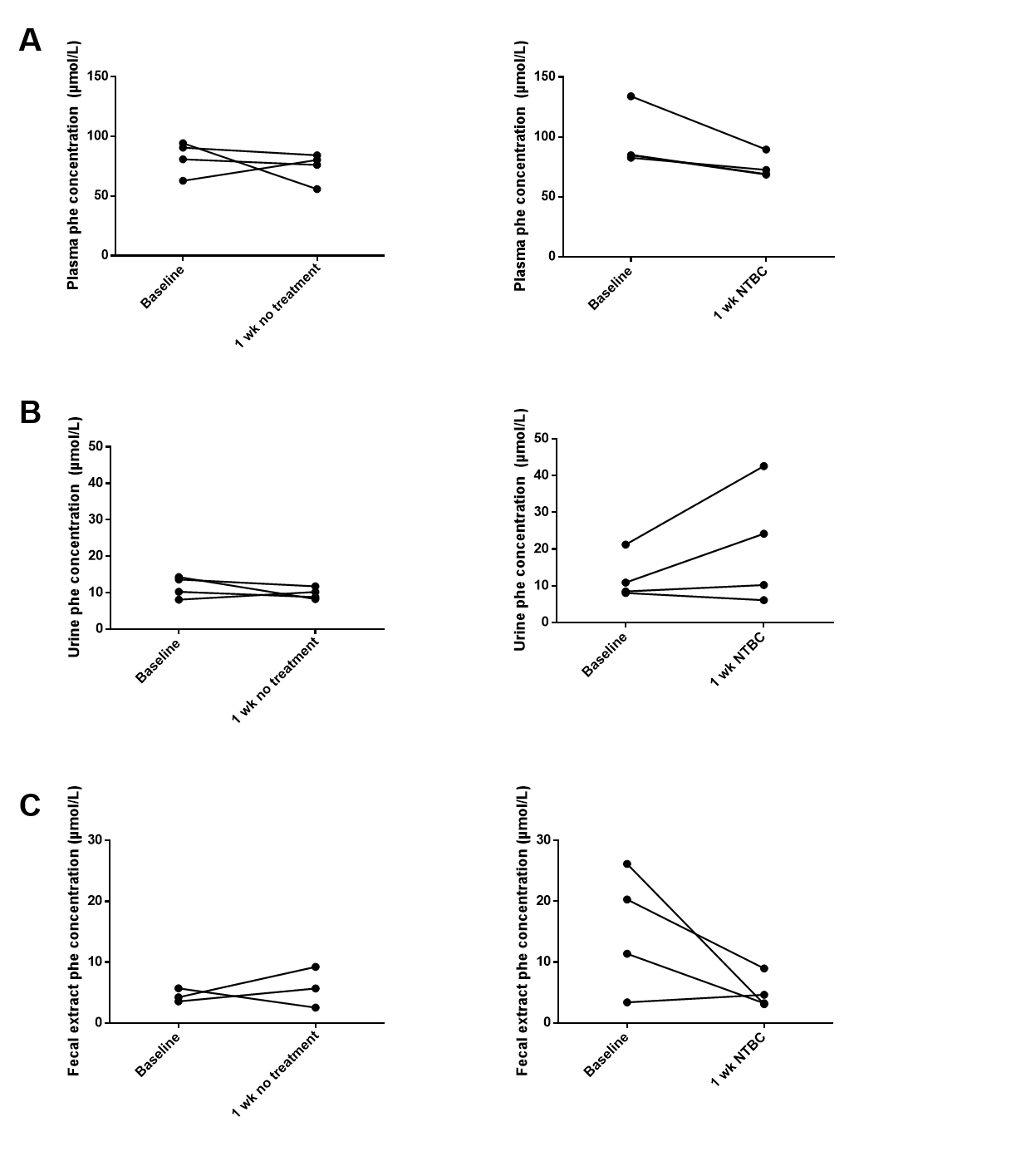

**Figure S3**. Phenylalanine concentrations at baseline then at one week on nitisinone (NTBC) or no treatment in plasma (A), urine (B) and faeces (C).

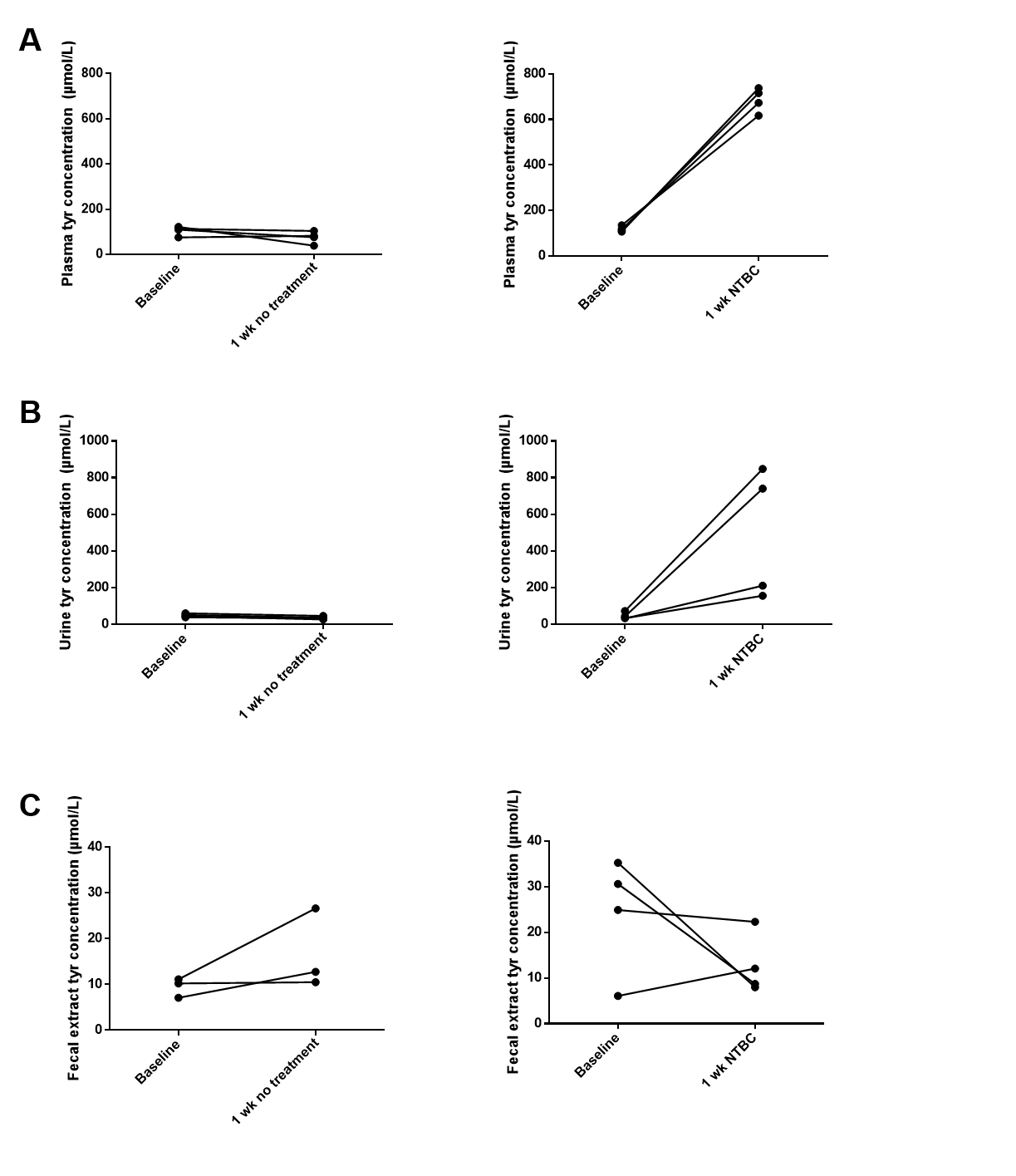

**Figure S4**. Tyrosine concentrations at baseline then at one week on nitisinone (NTBC) or no treatment in plasma (A), urine (B) and faeces (C).

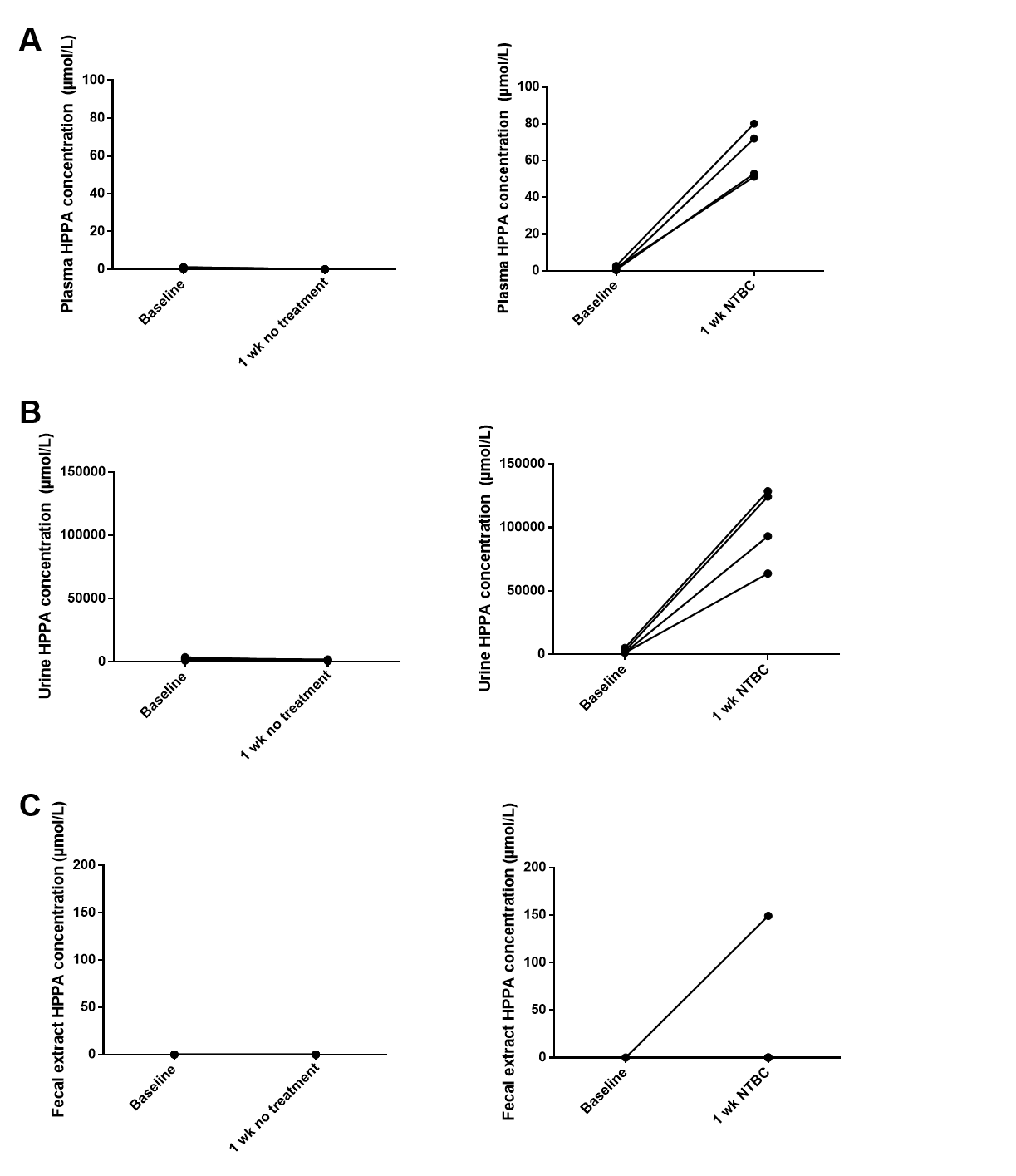

**Figure S5**. HPPA concentrations at baseline then at one week on nitisinone (NTBC) or no treatment in plasma (A), urine (B) and faeces (C).

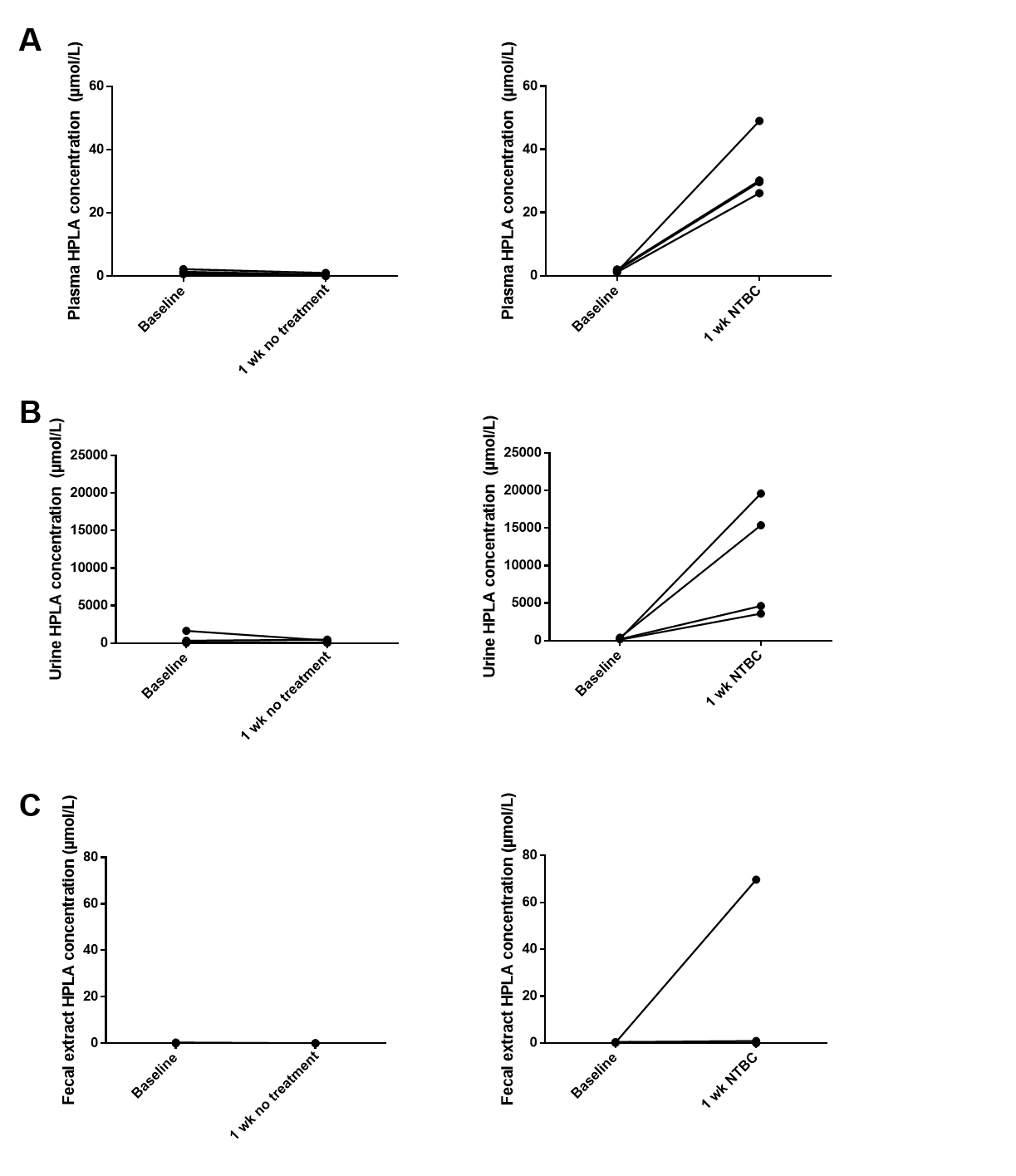

**Figure S6**. HPLA concentrations at baseline then at one week on nitisinone (NTBC) or no treatment in plasma (A), urine (B) and faeces (C).

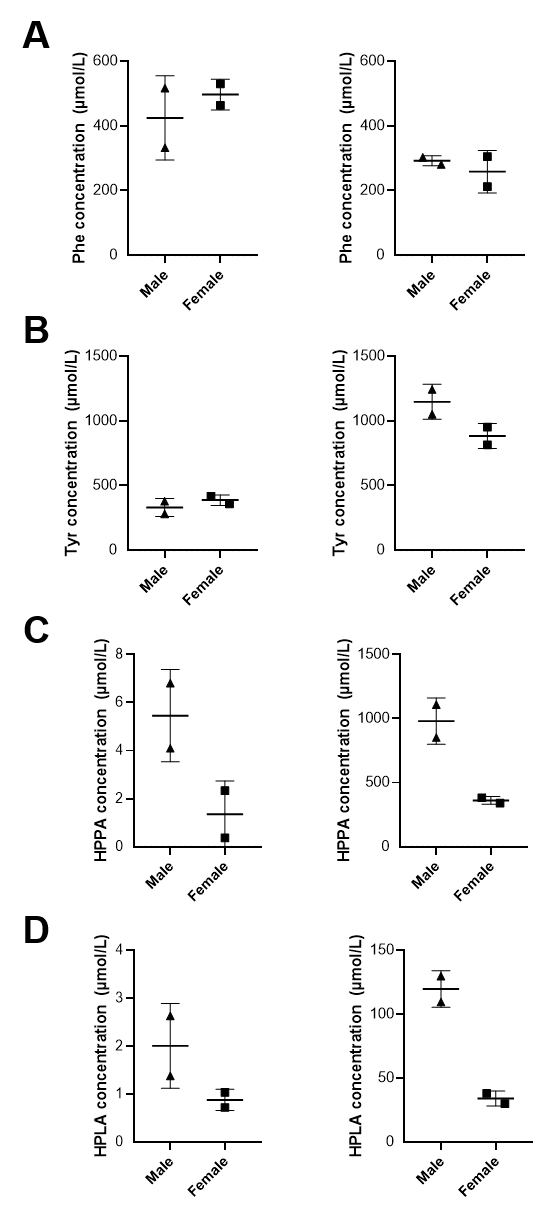

**Figure S7**. Comparison between male and female mice in bile metabolites phe (A), tyr (B), HPPA (C) and HPLA (D) at one week on nitisinone (right hand column) or no treatment (left hand column).
